# Selected reaction monitoring for the quantification of *Escherichia coli* ribosomal proteins

**DOI:** 10.1101/2020.07.16.206078

**Authors:** Yuishin Kosaka, Wataru Aoki, Megumi Mori, Shunsuke Aburaya, Yuta Ohtani, Hiroyoshi Minakuchi, Mitsuyoshi Ueda

## Abstract

Ribosomes are the sophisticated machinery that is responsible for protein synthesis in a cell. Recently, quantitative mass spectrometry (qMS) based on data-dependent acquisition (DDA) have been widely used to understand the biogenesis and function of ribosomes. However, DDA-based qMS sometimes does not provide the reproducible and quantitatively reliable analysis that is needed for high-throughput hypothesis testing. To overcome this problem, we developed a highly sensitive, specific, and accurate method to quantify all ribosomal proteins (r-proteins) by combining selected reaction monitoring (SRM) and isotope labeling. We optimized the SRM methods using purified ribosomes and *Escherichia coli* lysates, and verified this approach as a high-throughput analytical tool by detecting 41 of the 54 r-proteins separately synthesized in *E. coli* S30 extracts. The SRM methods will enable us to utilize qMS as a high-throughput hypothesis testing tool in the research of *E. coli* ribosomes, and they have potential to accelerate the understanding of ribosome biogenesis, function, and the development of engineered ribosomes with additional functions.

## Introduction

Ribosomes are the sophisticated machinery that synthesize proteins in all organisms. The *Escherichia coli* ribosome is a giant molecule composed of 30S and 50S subunits. The 30S subunit consists of 21 ribosomal proteins (r-proteins) and 16S ribosomal ribonucleic acid (rRNA), whereas the 50S subunit consists of 33 r-proteins, 5S rRNA, and 23S rRNA. The biogenesis of ribosomes is very complex, and several assembly-associated factors are involved in the process (1, 2). Studies of the function, structure, and assembly of ribosomes have been promoted in part by the reconstitution of the small and large subunits of ribosomes. The functional *E. coli* 30S subunit was first reconstituted by Traub & Nomura in 1968 (3). Due to the increased complexity of the structure, the reconstitution of the *E. coli* 50S subunit was not achieved until 1974 by Nierhaus & Dohme (4).

These early researchers revealed essential components of today’s knowledge about the assembly of ribosomes, which provides the basis of ribosome reconstruction and engineering. Jewett and coworkers (5-8) developed an *in vitro* integrated synthesis, assembly, and translation (iSAT) method that enables the co-activation of rRNA transcription, and the assembly of the transcribed rRNA with r-proteins into functional ribosomes in *E. coli* lysates. In contrast to the iSAT-based ribosome reconstruction, which uses a mixture of r-proteins, previous studies have also reported that individually purified ribosomal components can be reconstructed into 30S subunits (9-12). These approaches were integrated in a single reaction in the protein synthesis using recombinant elements (PURE) system (13), which coupled the transcription of 16S rRNA, assembly of the 30S subunit, and synthesis of sfGFP, which was translated by reconstructed ribosomes (14). Moreover, the autonomous synthesis and assembly of the 30S subunit on a chip using ribosomal genes was reported recently (15).

In parallel to these efforts on ribosome reconstruction, researchers have tried to redesign the translational machinery. Orthogonal translation systems based on modified Shine Dalgarno (SD)/anti-SD sequence pairs enable the translation of orthogonal mRNAs independent of the cellular translation system (16-19). This strategy has been applied to improving the site-specific incorporation efficiency of unnatural amino acids (20-23) and evaluating the activity of reconstructed or engineered ribosomes (14, 19, 24-26). In addition, tethered or stapled approaches (19, 24-27) have been established to facilitate the engineering of 50S subunits. Analyzing the structure-function correlation when mutations are introduced to the peptidyl transferase center (PTC) is also a target of ribosome engineering (28, 29), and a recent study established a high-throughput evaluation system using iSAT to evaluate comprehensive mutations in PTCs (30). The artificial evolution of 16S rRNA for the reconstruction of functional 30S subunits without post-transcriptional modifications is also achieved by combining iSAT, *in vitro* evolution, and a liposome sorting technique (31).

It is valuable to analyze the dynamics of each component to understand the process of ribosome reconstruction. For example, tracking expression levels or incorporation rates of each r-protein into ribosome assembly intermediates provides crucial information toward understanding the assembly process. Previous studies have revealed the composition of r-proteins and associated factors of ribosome assembly intermediates (2, 32, 33) and the stoichiometry of components of reconstructed ribosomes using mass spectrometry (12, 14, 26). However, conventional mass spectrometry based on data-dependent acquisition (DDA) may not be appropriate for the reproducible and reliable analyses that are required for high-throughput hypothesis testing. This problem can be overcome by conducting selected reaction monitoring (SRM), which enables the accurate and reproducible quantification of targeted proteins (34). The high-throughput and reproducible analysis of r-proteins using SRM can be a powerful method to accelerate the reconstruction and engineering of ribosomes. However, SRM methods have been developed for a limited number of r-proteins (35) and not for all 54 *E. coli* r-proteins.

In this study, we developed a highly sensitive, specific, and quantitatively accurate method to quantify all *E. coli* r-proteins by combining targeted proteomics using SRM with isotope labeling of nascent r-proteins. We determined optimized transitions for the quantification of 54 r-proteins using protein digests of purified ribosomes, *E. coli* lysate, and r-protein-overexpressed *E. coli* lysates. Further, we verified the SRM methods as a high-throughput analytical tool by detecting 41 of the 54 r-proteins separately synthesized in *E. coli* S30 extracts. This method will enable the utilization of quantitative mass spectrometry (qMS) as a high-throughput hypothesis testing tool in the field of ribosome research, and it has the potential to accelerate the understanding of ribosome biogenesis and development of functionally modified ribosomes (Fig 1).

**Fig 1.**
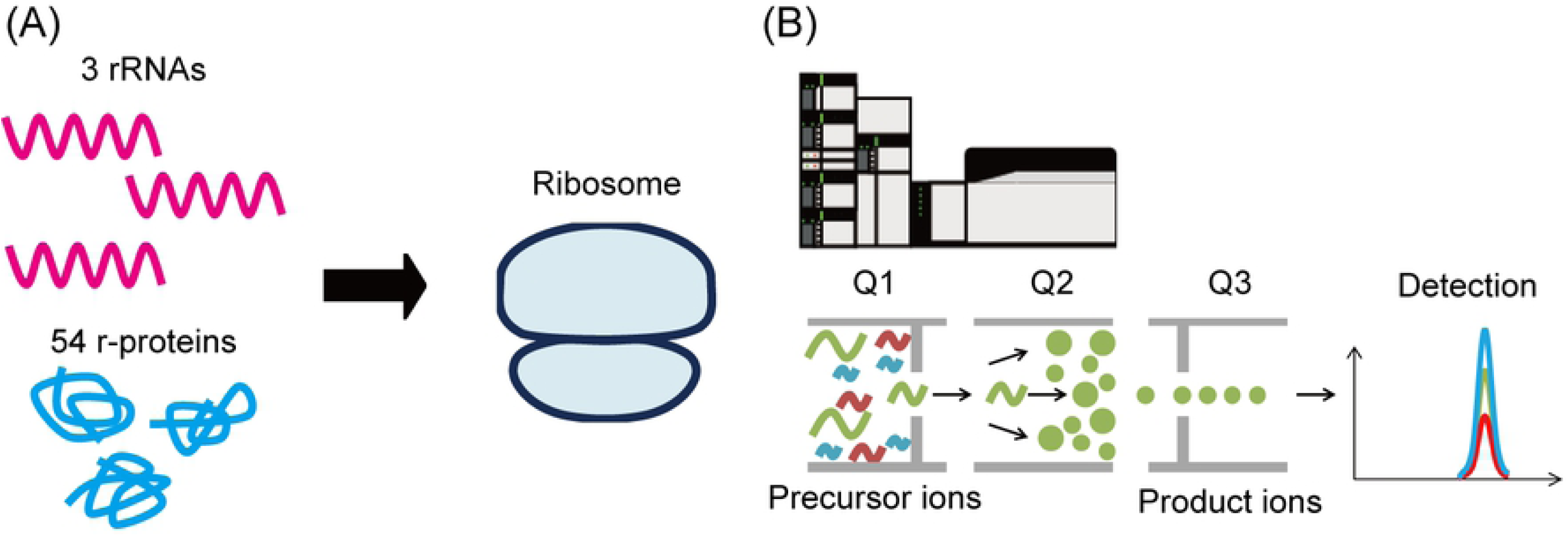
Selected reaction monitoring analysis for the quantification of *Escherichia coli* ribosomal proteins. (A) Biogenesis of *E. coli* ribosomes, including the assembly of three rRNAs and 54 r-proteins. (B) Nascent r-proteins and endogenous r-proteins were quantified using optimized SRM methods. The SRM methods enable us to utilize qMS as a high-throughput hypothesis testing tool for the research of *E. coli* ribosomes.

## Materials and methods

### Plasmids and strains

Genes encoding ribosomal proteins were amplified from the complete set of *E. coli* K-12 ORF archive (ASKA) library (36), and a vector backbone was amplified by PCR using pUC19 with KOD-Plus-Neo (TOYOBO, Osaka, Japan). The amplified fragments were gel-purified and fused using an In-Fusion^®^ HD Cloning Kit (Takara Bio, Shiga, Japan). Full sequences of the plasmids used in this study are provided in Supplementary Information I. In terms of the plasmids encoding each r-protein, we showed the sequence of pT7_rplA as a representative (Supplementary Information I).

The chromosomal *lacZ* gene of BL21 Star™ (DE3) (Thermo Fisher Scientific, Waltham, MA, USA) was replaced with the kanamycin-resistance gene (*kmr*) using Red-mediated recombination (37). The Red helper plasmid, pKD46, was transformed into the *E. coli* cells, and the transformants were cultured in 50 mL SOC medium with 100 μg/ml ampicillin (Viccillin^®^ for injection, Meiji Seika Pharma, Tokyo, Japan) and 10 mM L-(+)-arabinose (Nacalai Tesque, Kyoto, Japan). The fragment of the *kmr* gene was amplified using primers from pKD13: the H1P1 forward primer,5’-GAAATTGTGAGCGGATAACAATTTCACACAGGAAACAGCTGTGTAGGCTGGAGCTGCTTC-3’,andtheP4H2reverseprimer, 5’-TTACGCGAAATACGGGCAGACATGGCCTGCCCGGTTATTAATTCCGGGGATCCGTCGACC-3’. Then, it was purified using a MinElute^®^ PCR Purification Kit (QIAGEN, Hilden, Germany). Electroporation was performed using a Gene Pulser^®^ II (Bio-Rad, Hercules, CA, USA) with a Nepa Electroporation Cuvette with a 1 mm gap (Nepa Gene, Chiba, Japan) at 1350 V, 10 μF, and 600 Ohm. Next, the electroporated cells were incubated overnight at room temperature in SOC medium. After incubation, the cells were cultured on an LB agar plate containing 50 μg/ml kanamycin monosulfate (Nacalai Tesque) to select Km^R^ transformants. The deletion of the *lacZ* gene was verified using blue-white selection on an LB agar plate containing 0.1 mM isopropyl-β-D-thiogalactopyranoside (Nacalai Tesque) and 4.0 × 10^−3^ % 5-bromo-4-chloro-3-indolyl-β-D-galactoside (Takara Bio).

The Miraprep method (38) was performed using LaboPass™Plasmid Mini (Hokkaido System Science, Hokkaido, Japan) to prepare plasmids for cell-free protein synthesis (CFPS). Then, the extracted plasmids were purified further using a QIAquick^®^ PCR Purification Kit (QIAGEN).

### Preparation of S30 extract

S30 extract was prepared as previously reported (5-7). BL21 Star™(DE3) *lacZ::kmr* cells were grown in 1 L of 2×YPTG medium. Then, the cells were disrupted using an EmulsiFlex-C5 homogenizer (Avestin, Ottawa, Canada) with a single pass at a pressure of 20,000 psi. The lysate was centrifuged twice at 30,000 *g* for 30 min at 4 °C. Then, the supernatant was dialyzed using an iSAT buffer according to previous reports (7, 39). The protein concentration of the S30 extract was concentrated to 30–40 mg/mL, as determined using a Protein Assay BCA Kit (Nacalai Tesque).

### Cell-free protein synthesis

Cell-free protein synthesis was conducted according to previous reports with slight modifications (5, 8). The final concentration of T7 RNA polymerase (New England BioLabs, Ipswich, MA, USA) was 0.8 U/μL. Plasmid concentrations of pET41a_T7_sfGFP, pUC19_T7_LacZ, and the plasmids encoding ribosomal proteins were 80 nM, 60 nM, and 55.3 nM, respectively. The expression of sfGFP (40) from the pET41a_T7_sfGFP plasmid was induced using 2 mM of isopropyl-β-D-thiogalactopyranosides. CFPS solutions (15 μL) were incubated at 37 °C for 3 h in a black-walled, polystyrene 96-well plate with a solid bottom and half volume (Greiner Bio-One International GmbH, Kremsmünster, Austria). The expression of sfGFP and LacZ was monitored using a Fluoroskan Ascent FL™ 96-well-plate reader (Thermo Fisher Scientific) at λ_ex_ = 485 nm and λ_em_ = 510 nm. To quantify the activity of lacZ, 3.3 nM of 5-chloromethylfluoresecein di-beta-D-galactopyranoside (CMFDG) (Invitrogen, Waltham, MA, USA) was added to the reaction solution. For isotope labeling, we used 20 amino acids that contained 2 mM of ^13^C_6_ ^15^N_2_ L-lysine (Thermo Fisher Scientific) and ^13^C_6_ ^15^N_4_ L-arginine (Thermo Fisher Scientific) instead of ^12^C_6_ ^14^N_2_ L-lysine and ^12^C_6_ ^14^N_4_ L-arginine (Sigma-Aldrich Corporation, St. Louis, MO, USA). The synthesized proteins were quantified using a triple quadrupole liquid chromatograph mass spectrometer LCMS-8060 (Shimadzu, Kyoto, Japan), as described below.

### Overexpression of r-proteins in *Escherichia coli*

The plasmids encoding each ribosomal protein were purified using the ASKA library (36). Then, the plasmids were introduced into *E. coli* BL21 (DE3) *lacZ::kmr*. The cells were grown at 37 °C to OD_600_ = 0.5–0.6 in 5 mL LB medium containing 100 μg/ml ampicillin (Viccillin^®^ for injection, Meiji Seika Pharma), incubated at 37 °C for 3 h with 0.1 or 1 mM IPTG (Nacalai tesque), and then centrifuged at 3,000 rpm for 5 min. Then, the pellets of the cells were suspended in 200 μL of lysis buffer of 2 M urea, 50 mM ammonium bicarbonate, and with a pH of 8.0 and transferred into 1.5 mL microtubes in an ice-water bath. The cells were disrupted using a sonicator (BIORUPTOR UCD-250, Cosmo Bio, Tokyo, Japan) for 10 min at 4 °C (Level 5, a 30-s sonication with 30-s interval). Then, the lysates were centrifuged at 14,000 g for 10 min at 4 °C, and the supernatants were collected. The ribosomal proteins were quantified using the LCMS-8060 (Shimadzu, Kyoto, Japan) as described below.

### Preparation of protein digests

The protein samples of purified ribosome (New England BioLabs), *E. coli* lysate, *E. coli* lysates containing each overexpressed r-protein, r-proteins produced in S30 extract, and LacZ produced in S30 extract were mixed with 120 μL of methanol, 30 μL of chloroform, and 90 μL of ultrapure water, which were added to 30 μL of each protein sample and vortexed thoroughly. Then, the mixtures were centrifuged at 13,000 *g* for 5 min. The top aqueous layer was pipetted off, and 90 μL of methanol was added to the solution. Then, the solution was mixed thoroughly and centrifuged at 13,000 *g* for 5 min. The aqueous layer was pipetted off and the pellet was air-dried for 5 min. Then, the pellet was dissolved in a lysis buffer of 2 M urea and 50 mM ammonium bicarbonate with a pH of 8.0. After alkylation and reduction steps (41), proteolytic enzymes, trypsin (Promega, Madison, WI, USA) and/or lysyl endopeptidase (FUJIFILM Wako Pure Chemical Corporation, Osaka, Japan) were added to the samples at the ratio of total proteins/proteolytic enzyme = 75:1 (w/w). The samples were shaken at 37 °C for 24 h, and then the reaction was stopped by adding trifluoroacetic acid to provide a final concentration of 1% (v/v). The digested peptides were desalted using a MonoSpin^®^ C18 column (GL Sciences, Tokyo, Japan) and dried using vacuum centrifugation. Then, the dried samples were eluted to 5 μg/μL using 10% formic acid. Next, the peptide samples were filtered through 0.45 μm filters (Merck KGaA, Darmstadt, Germany). For the digestion of rplF, rplT, rplY, and rpmC that were synthesized in S30 extracts, samples were digested using Lys-C for 4 h, and then they were digested using trypsin for 24 h.

### Quantification of ribosomal proteins using triple quadrupole liquid chromatography-mass spectrometry

The digested proteins were analyzed using an UltiMate 3000 RSLCnano (Thermo Fisher Scientific) and an LCMS-8060 (Shimadzu, Kyoto, Japan) triple quadrupole mass spectrometer. A 500 mm monolithic column with a100 μm ID (42) was connected to a six-port injection/switching valve (Valco Instruments, Houston, TX, USA). A 5 μm, 0.3×5 mm L-column ODS (Chemical Evaluation and Research Institute, Saitama, Japan) was used as a trap column. The samples were passed through a 20 μL sample loop and injected into the monolithic column at a flow rate of 500 nL/min. Then, a gradient was generated by varying the mixing ratio of the two eluents: 0.1 % (v/v) formic acid diluted with ultrapure water (eluent A); and 0.1 % (v/v) formic acid diluted with acetonitrile (eluent B). The gradient started at 5% of eluent B over 7.5 min, it was then increased to 40% of eluent B for 27.5 min, and it was finally increased to 95% of eluent B for a 5-min hold. Then, the ratio of eluent B was quickly adjusted to the initial composition and held for 5 min to re-equilibrate the column. The column oven was kept at 40 °C, and the autosampler was kept at 4 °C. The block and the interface temperature were adjusted to 400 °C and 300 °C, respectively. Using Skyline software (43), we prepared SRM methods for both non-labeled proteins and proteins that were labeled with ^13^C_6_ ^15^N_2_ L-lysine and ^13^C_6_ ^15^N_4_ L-arginine. Double-charged and triple-charged states of 6–25 mer peptides were selected to predict precursor ion candidates. Peptides, including methionine, were excluded from the candidates because they are susceptible to oxidation and unsuitable for quantitative analysis. Carbamidomethyl cysteine was set as a fixed modification. For fragment ions, singly charged or doubly charged b- or y-series ions with 50–1500 m/z were predicted. All selected peptides were evaluated for the uniqueness of their sequences using an *E. coli* background, and the SRM methods were scheduled based on the obtained retention time with a dwell time of 5 ms.

### Data analysis

We analyzed protein digests of purified ribosome (New England BioLabs) and purified LacZ (FUJIFILM Wako Pure Chemical Corporation) and drew calibration curves to quantify the r-proteins and LacZ synthesized in S30 extracts. Statistical analysis was conducted using a two-tailed *t*-test. We calculated the ratio of peak areas from isotope-labeled ‘heavy’ r-proteins and ‘light’ endogenous r-proteins to compare the abundance ratio of r-proteins synthesized in S30 extracts to the endogenous counterparts. The names of r-proteins were written according to the universal nomenclature for r-proteins (44). The datasets generated and/or analyzed during the current study are available in the jPOST repository (accession number: JPST000907 or PXD020266) (45).

## Results

### Development of selected reaction monitoring methods for the quantification of r-proteins

For the SRM analysis, we selected peptides that met the following criteria: (1) peptides that show intense peaks when prepared using purified ribosomes, (2) peptides that show intense peaks when prepared using *E. coli* lysate and show the same retention times observed at (1), and (3) peptides that show stronger intensities when prepared using r-protein-overexpressed *E. coli* lysates compared with that of (2). We attempted to select three peptides for each r-protein and 3–4 fragment ions for each peptide to enable precise quantification.

We predicted precursor-to-fragment ion transitions using Skyline for each protein of interest, analyzed the protein digests prepared using purified ribosomes, and selected candidate transitions for each r-protein (S1 Fig). The selected transitions were then verified by analyzing the protein digests prepared using *E. coli* lysate or r-protein-overexpressed *E. coli* lysates, as described in the Materials and methods section (S1 Fig). We successfully selected specific transitions for each r-protein, as seen in S2 Fig, and the optimized SRM methods are summarized in S1 Table. The linearity of the SRM methods was determined using a dilution series of protein digests of purified ribosomes. A dilution rate of 1.0 represents a 5 μL injection of 8 μg/mL of the samples. The peak area of each peptide was plotted against its concentration, and most transitions showed a high correlation coefficient (S2 Table).

### Evaluation of cell-free protein synthesis using S30 extracts

We attempted to quantify r-proteins separately produced in *E. coli* S30 extracts using our SRM methods as a high-throughput analytical tool. To evaluate the activity of the *E. coli* S30 extracts, we produced sfGFP in the *E. coli* S30 extract, and achieved a reaction yield of about 2.0 mg/mL **(S3 Fig**), which is comparable to that of previous reports (6, 39). To evaluate the production level of r-proteins, we discriminated nascent r-proteins from endogenous r-proteins. Then, we planned to incorporate isotope-labeled amino acids into the nascent r-proteins. However, endogenous amino acids in S30 extracts can also be incorporated into nascent r-proteins. Therefore, we determined the incorporation rates of endogenous and exogenous amino acids into the nascent proteins. As a model, we attempted to quantify the incorporation rates of isotope-labeled amino acids into the LacZ that was synthesized in S30 extracts derived from the *E. coli* BL21 Star™(DE3) *lacZ::kmr* strain. First, we developed SRM methods for LacZ with high specificity and accuracy (S4 Fig). Then, we synthesized LacZ in the S30 extracts with or without isotope-labeled amino acids, and we determined peptide ratios of heavy to light (Fig 2). Consequently, we found that most of the nascent LacZ consisted of only isotope-labeled amino acids. This is because the cell lysates were dialyzed using an iSAT buffer, which washed out almost all of the endogenous amino acids. These results suggest that the intensity of the ‘heavy’ peptides directly corresponds to the protein production level in the S30 extracts that were prepared in this study.

**Fig 2.**
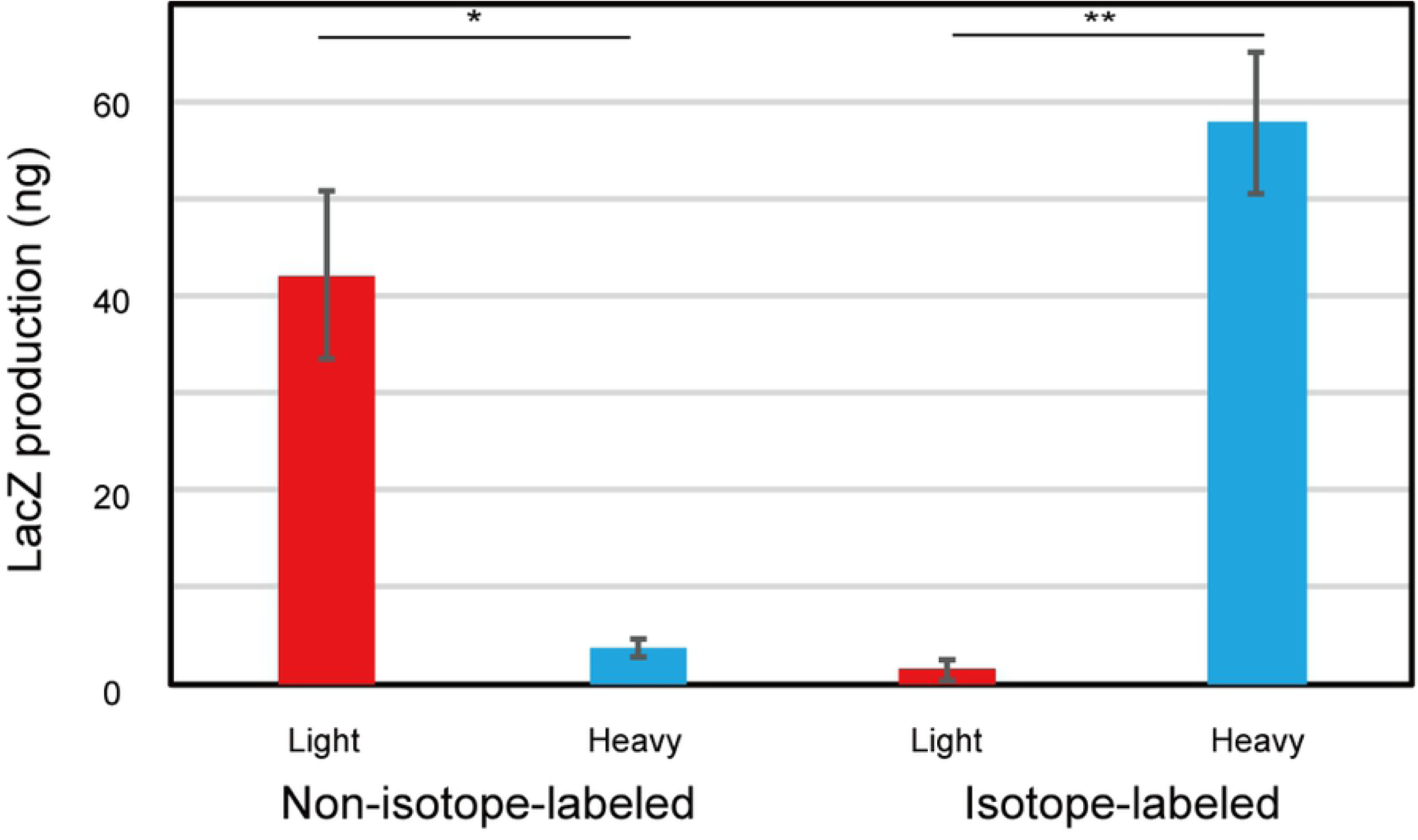
Quantification of LacZ synthesized in S30 extract. Quantification of LacZ synthesized with or without 2 mM of ^13^C_6_ ^15^N_2_ L-lysine and ^13^C_6_ ^15^N_4_ L-arginine in S30 extracts. The protein digests were quantified using the selected reaction monitoring methods for LacZ. The protein yield was deduced based on the peak area of the transitions. A non-isotope-labeled control was used to measure background signals. Data are shown as mean ± standard error of the mean (N = 3). Statistical significance was determined using a two-tailed Student’s *t*-test with significance at *p <0.05, **p <0.01. The blue bar indicates heavy proteins and the red bar indicates light proteins.

### Profilingofnascentr-proteinsseparatelyproducedinS30 extracts

We quantified isotope-labeled nascent r-proteins that were separately synthesized in S30 extracts using our SRM methods. We synthesized each r-protein in S30 extracts containing 2 mM of ^13^C_6_ ^15^N_2_ L-lysine and ^13^C_6_ ^15^N_4_ L-arginine instead of ^12^C_6_ ^14^N_2_ L-lysine and ^12^C_6_ ^14^N_4_ L-arginine. The S30 extracts were digested and analyzed as described in the Materials and methods section, and the ratios of nascent r-proteins (‘heavy’) to endogenous r-proteins (‘light’) were calculated (Heavy/Light ratio) to investigate the abundance ratio of nascent proteins to endogenous counterparts. The r-proteins, which had an S/N ratio >5 were defined as detected proteins (Fig 3). As a result, 41 of the 54 ‘heavy’ r-proteins were successfully detected while the others were undetected: bL21, bL27, uL30, bL31, bL34, bL35, uS2, uS7, uS8, uS13, bS16, uS19, and bS21. Although several r-proteins showed higher intensity peaks, many of the detected r-proteins had lower intensity peaks than their endogenous counterparts.

**Fig 3.**
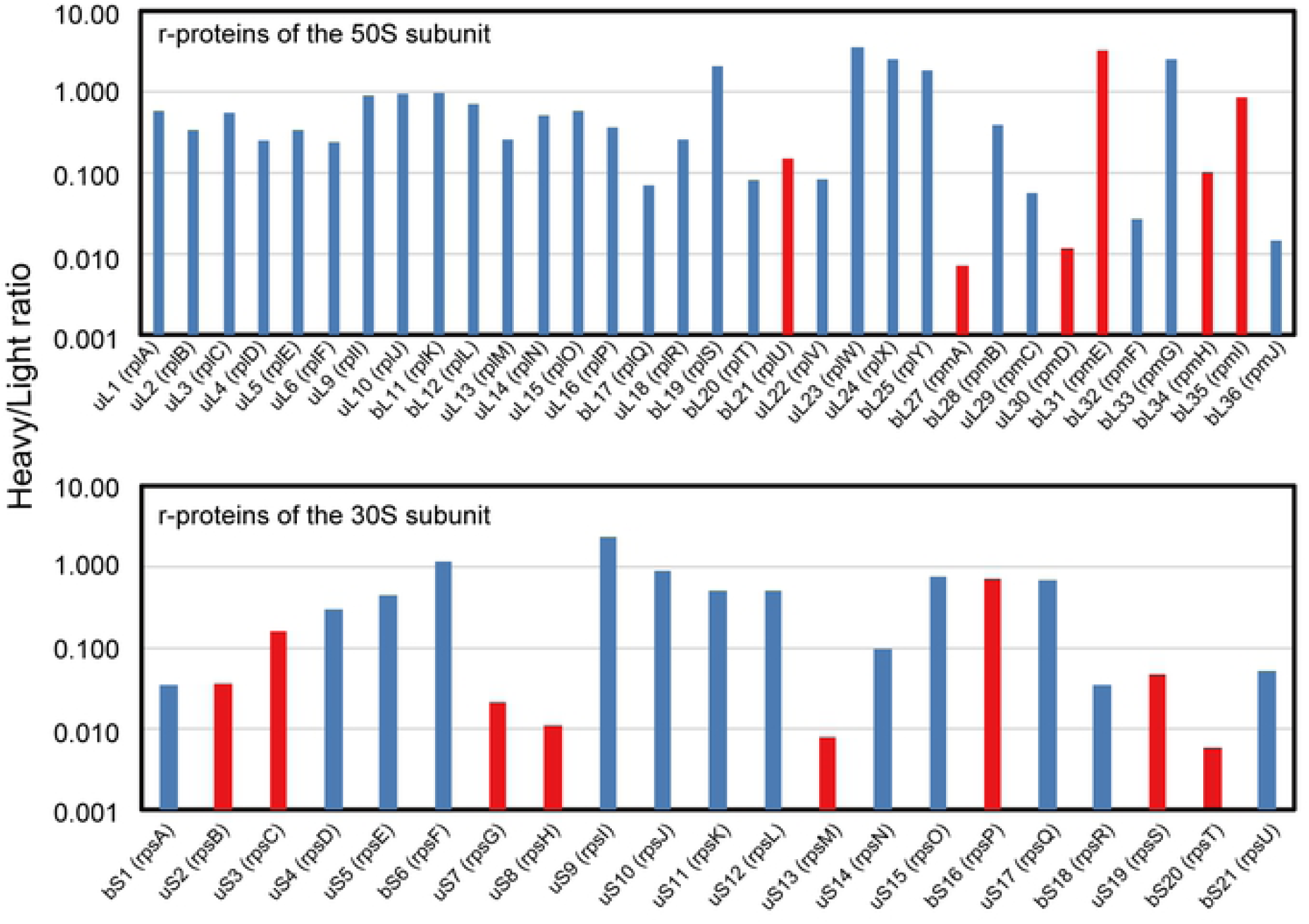
Quantification of r-proteins synthesized in S30 extracts. Isotope-labeled r-proteins were separately synthesized in S30 extracts and digested using trypsin or lysyl endopeptidase. The digests were quantified using optimized selected reaction monitoring methods. The ratios of nascent r-proteins (‘heavy’) to endogenous r-proteins (‘light’) were calculated (Heavy/Light ratio). The r-proteins with a S/N ratio >5 were defined as detected proteins. The blue bar indicates detected proteins and the red bar indicates undetected proteins.

## Discussion

Quantitative mass spectrometry based on DDA, which can detect proteins with high sensitivity, has been an effective technology for understanding ribosome biogenesis and the development of functionally modified ribosomes. Data-dependent acquisition-based qMS was used to measure relative production levels of r-proteins (46), and more recently, has revealed the composition of r-proteins and associated factors of ribosome assembly intermediates (2, 32, 33). A flexible standard has also been developed using QconCAT technology to capture the composition of stable or transient ribosome complexes using orbitrap and MALDI-TOF mass spectrometry (47). Furthermore, the DDA approach has been used for analyzing *in vitro* ribosome reconstruction (12, 14) and ribosome engineering (48).

However, in DDA-based qMS, the acquisition of MS/MS spectra can be stochastic and DDA-based qMS does not always detect all of the target peptides when samples contain a large number of proteins or peptides can be co-eluted (49). Therefore, DDA may not be appropriate for conducting reproducible and reliable analyses, which are required for high-throughput hypothesis testing (50). Selected reaction monitoring using triple quadrupole MS can detect and quantify specific ions by setting a combination of m/z of a first quadrupole (Q1) filter that passes precursor ions and a third quadrupole (Q3) filter that passes fragment ions after collision-induced dissociation. Normally, multiple transitions (Q3) are set for a single peptide (Q1). Hence, these transitions are detected as co-eluting peaks in a chromatogram, which can guarantee their specificity (34, 51-53)

So far, SRM methods have been developed against the limited number of r-proteins (35), and not against all 54 *E. coli* r-proteins. In this study, we developed a method to quantify all of the *E. coli* r-proteins with high sensitivity, specificity, and quantitative accuracy by combining SRM with isotope labeling of nascent r-proteins. We determined optimized transitions for the quantification of 54 r-proteins using protein digests of purified ribosomes, *E. coli* lysate, and r-protein-overexpressed *E. coli* lysates (S1 Table). The quantification of isotope-labeled LacZ synthesized in S30 extracts derived from an *E. coli lacZ::kmr* strain confirmed that nascent proteins were specifically labeled by isotope amino acids, which enabled precise quantification of nascent proteins (Fig 2). We verified the SRM approach developed in this study as a high-throughput analytical tool by quantifying all of the r-proteins separately synthesized in *E. coli* S30 extracts.

Some of the r-proteins were not detected when we separately expressed them in S30 extracts: bL21, bL27, uL30, bL31, bL34, bL35, uS2, uS7, uS8, uS13, bS16, uS19, and bS21. These r-proteins included r-proteins smaller than 9 kDa, such as uS2, uS7, uS8, and uS13, which are assumed to be difficult to use for selecting peptides for SRM partly because only a limited number of peptide candidates can be generated from small r-proteins. This problem can be resolved by using proteases other than trypsin or lysyl endopeptidase. In addition, bL17, bL20, bL29, bL36, bS1, and bS21 showed a low expression level, as reported previously (12). The inefficient expression of some r-proteins can be improved by increasing the translation and protein folding efficiency in S30 extracts by using optimized plasmid design (54), polycistronic expression (55) and the macromolecular crowding effect (56). Moreover, a pipet-tip-based peptide micropurification system, which enables the multidimensional fractionation, desalting, filtering, and concentration (57) of peptides from reacted S30 extracts, might improve the detection sensitivity of these peptides.

In conclusion, we developed a highly sensitive, specific, and accurate high-throughput quantification method of 54 *E. coli* r-proteins by combining SRM with isotope labeling of nascent r-proteins. The SRM methods for r-proteins established in this study enable the profiling of ribosome biogenesis by quantifying each r-protein in a highly quantitative manner, and they enable us to utilize qMS as a high-throughput hypothesis testing tool in the field of ribosome research. This study provides a powerful method for understanding ribosome biogenesis, and it accelerates research on the reconstruction or engineering of *E. coli* ribosomes.

## Acknowledgments

We thank the National BioResource Project (NBRP: *E. coli*) for providing the ASKA library. W.A. was supported by the Japan Society for the Promotion of Science (grant No. 19K16109 and 26830139; https://www.jsps.go.jp/english/) and Sugiyama Chemical & Industrial Laboratory (http://www.sugiyama-c-i-l.or.jp/). The funders had no role in study design, data collection and analysis, decision to publish, or preparation of the manuscript.

## Supporting Information

**S1 Fig. Ion chromatograms corresponding to peptides derived from r-proteins**. Candidate peptides for selected reaction monitoring analysis were selected using Skyline software. We quantified the prepared r-proteins using purified ribosomes, *E. coli* lysate, and r-protein-overexpressed *E. coli* lysates using triple quadrupole liquid chromatography-mass spectrometry.

**S2 Fig. Ion chromatograms corresponding to the selected transitions associated with peptides from ribosomal proteins**. For each peptide, several transitions with intense peaks were selected based on the quantification of r-proteins from purified ribosomes. The calibration curves of all transitions are described in Table S2.

**S3 Fig. Validation of the protein production activity of S30 extracts**. sfGFP was expressed in S30 extracts prepared using an EmulsiFlex-C5 homogenizer. The expression of sfGFP was monitored using a 96-well-plate reader at λ_ex_ = 485 nm and λ_em_ = 510 nm. Data are shown as the mean ± standard error of the mean (N = 3).

**S4 Fig. Development of selected reaction monitoring methods for LacZ**

(A) An ion chromatogram corresponding to the transitions associated with peptides from LacZ. Tryptic digests of LacZ standard were analyzed by triple quadrupole liquid chromatography-mass spectrometry.

(B) Ion chromatograms corresponding to the selected transitions associated with peptides from LacZ. Tryptic digests of LacZ standard were quantified using triple quadrupole LC-MS/MS, and transitions with intense peak were selected.

(C) Calibration curves of transitions for the quantification of LacZ. Transitions that provide the highest R^2^ value were selected as the transitions with high accuracy. The dilution rate of 1.0 corresponds to the 5 μL injection of 1 μg/μL of the tryptic LacZ digest.

**S1 Table. Selected reaction monitoring methods for r-proteins**

**S2 Table. Calibration curves of selected reaction monitoring methods for r-proteins**

**S3 Table. Selected reaction monitoring methods for LacZ**

